# An Efficient Graph Algorithm for Diploid Local Ancestry Inference

**DOI:** 10.1101/2025.07.05.662656

**Authors:** Nafiseh Jafarzadeh, Jordan M Eizenga, Benedict Paten

## Abstract

In this paper, we present diploid sequence graphs, graphs whose paths encode pairs of haplotypes. We describe an efficient algorithm for creating a diploid graph from a directed acyclic (haploid) sequence graph, such that the diploid graph represents all the possible pairings of haplotypes present in the sequence graph and their similarity relationships.

Starting with the sequence graph, our method uses a graph decomposition approach based on an extension of the SPQR-tree to systematically identify structural patterns that reduce redundancy while preserving genetic variation. We develop a polynomial-time algorithm that parsimoniously enumerates all disjoint paths with shared endpoints in two-terminal directed acyclic graphs.

In the future, we envisage that diploid graphs may enable more accurate modeling of recombination, phasing, and variation-aware alignment in diploid genomes.

## Introduction

Sequence graphs provide a unified framework for modeling all types of sequence variation [1]. They are being used as a basis for building comprehensive pangenomes [2]. Efficient methods for a variety of genomic tasks have been developed and continue to be improved to use sequence graphs [1]. This includes mapping algorithms for mapping sequences to sequence graphs [3, 4], as well as haplotype inference algorithms [5]. Such algorithms reduce bias relative to using algorithms that instead use a linear reference representation.

Sequence graphs can encode haplotypes as walks through the graph, with the haplotype sequence given by concatenating the sequences of the vertices. This makes sequence graphs useful for reasoning about individual haplotypes. However, in humans and many other species, genomes are diploid: they have two copies of each chromosome. To date, there has been little attention to graph models that explicitly represent diploid genomes. Here we consider this challenge, and in particular, how to convert an input sequence graph into a diploid representation in which walks encode diploid combinations of genomes. Given such a representation, we show that it is possible to reason efficiently about recombination between pairs of diploid haplotypes.

### 1 Diploid Graph Modeling

In this section, we present a conceptual algorithm for constructing a diploid sequence graph that encodes all possible pairings of haplotype paths in an input sequence graph, assuming we already know the locations of the graph’s *hetero-positions*, which are decomposable local graph structures that we define below. A sequence graph is a graph where each vertex is labeled with a nucleotide string, and directed edges connect vertices based on relationships within the haplotypes. While more expressive representations are possible (i.e. bidirected graphs [6, 7]), here we assume for simplicity that sequence graphs are two-terminal directed acyclic graphs (DAGs), where maximal paths from the source to the sink correspond to haplotype sequences. In contrast, the vertices of the diploid-sequence graph are labeled with pairs of nucleotide strings, and maximal paths correspond to diplotype sequences.

Given a sequence graph with *n* maximal paths, a naive diploid construction pairs all distinct haplotypes, yielding 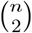 combinations. Since *n* often grows exponentially with the size of the graph, this results in a diploid graph that is both large and highly redundant. Moreover, by capturing only simple pairwise combinations, it eliminates the sequence graph’s implicit homology information and overlooks richer patterns of haplotype variation.

In contrast, our method builds on the naive approach but improves it by pairing only disjoint segments of haplotypes. Specifically, we pair regions corresponding to heterozygous positions, while representing homozygous regions only once. This strategy keeps the graph compact, preserves diploid-level topology, and captures the underlying structure of variation.

Building on this model, we propose a three-step approach for constructing a diploid sequence graph. First, we create a subgraph representing homozygous diploids, called the *homo-sequence graph*. This graph is isomorphic to the sequence graph, with each vertex labeled as (*v, v*), representing *n* homozygous diploids for *n* haplotypes. Figure 1 shows the sequence graph (A) and the homo-sequence graph (B).

**Figure 1.**
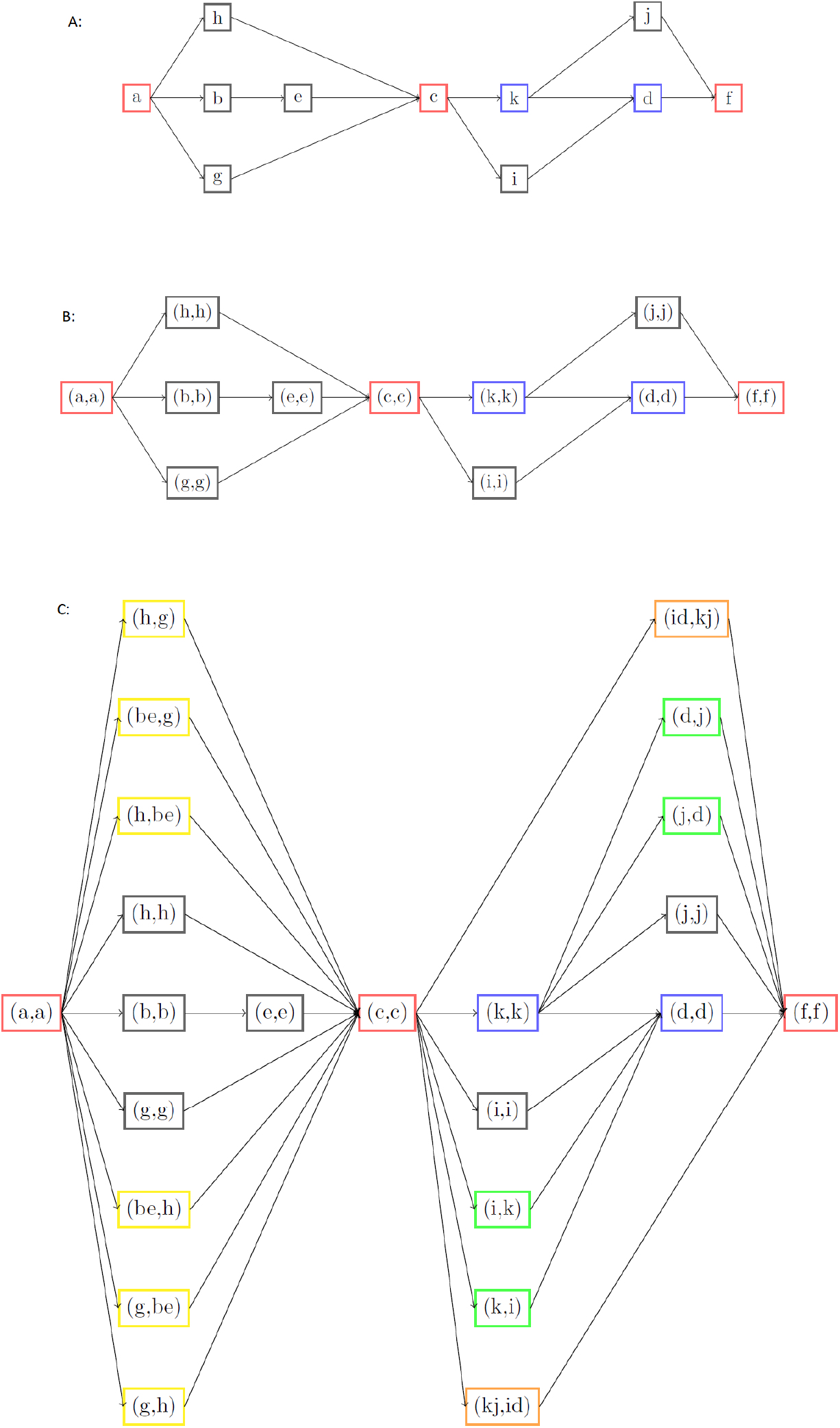
(A) A sequence graph. (B) The homo-sequence graph of (A). (C) The diploid-sequence graph constructed by extending (B) at hetero-positions.

Next, to add heterozygous diploids, we identify specific pairs of vertices in the sequence graph, which we refer to as hetero-positions. A hetero-position is defined as a pair of vertices like (*x, y*) where *x < y*, and which admit two paths between them such that the paths are disjoint except at *x* and *y*. Intuitively, hetero-positions mark the boundaries of variation between alternative haplotype paths in the graph.

In Figure 1, you can see all hetero-positions in (B) as four pairs of vertices in red and blue: ((*a, a*), (*c, c*)), ((*c, c*), (*f, f*)), ((*c, c*), (*d, d*)) and ((*k, k*), (*f, f*)).

Finally, we extend the graph by adding new vertices between hetero-positions to represent disjoint path pairs. In the same figure (C), new vertices in yellow, green, and orange are added between hetero-positions.

In Algorithm 1, we describe a generic procedure for constructing a diploid sequence graph under the assumption that the hetero-positions and their corresponding disjoint path pairs are given in advance.

Proposition 1 establishes the correctness of this algorithm.

#### Algorithm 1

Generic Diploid Sequence Graph Construction

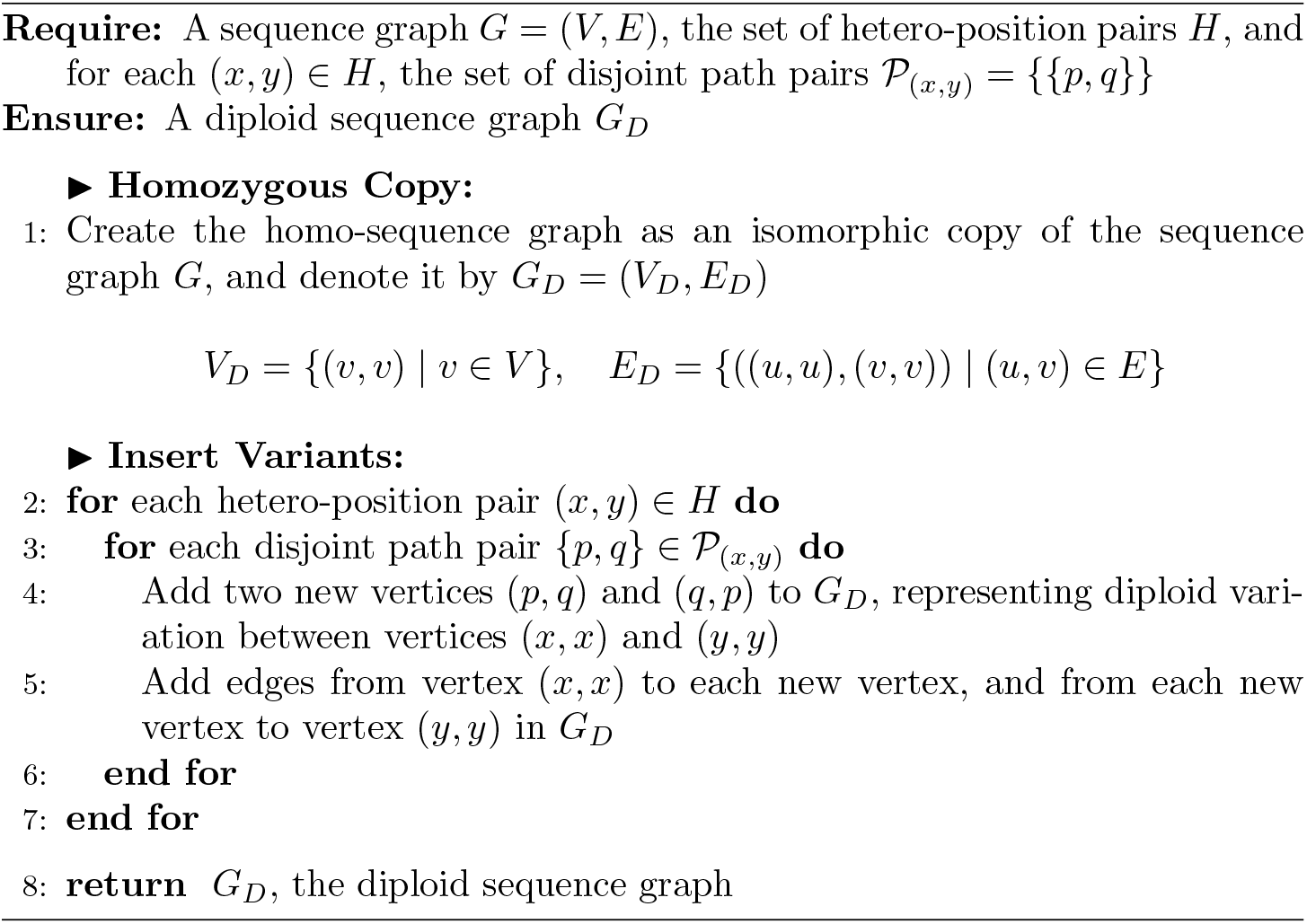

#### Proposition 1

(Validity of Algorithm 1). *Given a sequence graph G, a set of hetero-positions, and corresponding disjoint path pairs between each hetero-position, Algorithm 1 constructs a diploid sequence graph G*_*D*_ *that encodes all intended diploid sequences as maximal paths*.

*Proof*. The algorithm begins by creating an isomorphic copy of the sequence graph *G*, ensuring that all homozygous diploid sequences are preserved in *G*_*D*_. Now, consider two maximal paths *P* and *Q* in *G*, representing two haplotypes. We divide these paths into identical and distinct segments. The identical segments are present in *G*_*D*_ by construction. Each pair of distinct segments is bounded by a hetero-position pair (*x, y*) *∈ H*. Since disjoint path pairs like (*p, q*) and (*q, p*) are explicitly given for the hetero-position pair (*x, y*), and the algorithm connects them between vertices (*x, x*) and (*y, y*), the distinct segments of *P* and *Q* are also realized in *G*_*D*_. Therefore, the haplotype pairs (*P, Q*) and (*Q, P*), including both their identical and distinct segments, is represented as a maximal diploid path in *G*_*D*_.

Figure 1 shows a small example illustrating our method, from the original sequence graph to the homo-sequence graph and then to the diploid graph. The sequence graph in (A) contains 9 maximal paths. The naive method pairs all of them, creating 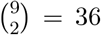 new vertices. In contrast, our method only pairs disjoint segments of paths, adding just 12 new vertices. This keeps the graph much smaller while still representing all diploid sequences.

### 2 Structural Decomposition

To identify hetero-positions and perform extensions, we need to enumerate all pairs of disjoint paths that share the same start and end vertices in a DAG. To do this, we propose a method based on a decomposition technique using the SPQR-tree framework [8].

The SPQR-tree is a data structure that decomposes a biconnected graph into a hierarchical representation of its structural components. It classifies the graph into four types of nodes. An *S*-node represents a series composition, where edges form a path-like structure. A *P* -node captures parallel connections, where multiple edges exist between the same pair of vertices. A *Q*-node corresponds to a single edge in the original graph. An *R*-node represents a triconnected subgraph that cannot be further decomposed into series or parallel structures. A *split pair* is a pair of vertices whose removal separates a graph component from the rest of the graph and serves as a connection point in the decomposition. Each internal node in the SPQR-tree is associated with a *skeleton graph*, a simplified structure built around a split pair. The skeleton graph includes a *virtual edge* in place of every subcomponent that would result from removing the split pair, capturing how subcomponents are connected. For *P* -nodes and *S*-nodes, the skeleton graph consists of multiple parallel edges or a cycle, respectively.

Let *G* be a sequence graph with source *s* and sink *t*. As a foundation for our approach, we aim to construct an SPQR-tree that reflects the DAG structure of *G*. Since SPQR-trees are defined for undirected biconnected graphs, we first convert *G* into an undirected graph and add a back edge (*t, s*) to ensure biconnectivity, which is necessary for the SPQR tree to be well-defined. We denote this undirected biconnected graph as 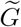. We then build the SPQR-tree of 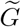 using the linear-time algorithm from [9], rooting it at the Q-node corresponding to the added edge (*t, s*). This rooting defines a consistent hierarchy and assigns a unique outer split pair (*x, y*) to each internal node. The outer split pair of a node is given by the endpoints of the virtual edge in its skeleton that connects it to its parent in the tree. These pairs determine the attachment points of each component and guide how subgraphs are recursively composed.

Finally, we restore directions using the original orientation of edges in *G*. We show that each *S*-node and *P* -node defines a two-terminal subgraph in the directed setting.

#### Lemma 1.

*Each S-node and P-node in the SPQR-tree of* 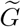, *rooted at the Q-node for* (*t, s*), *defines a two-terminal subgraph in the original G, with terminals equal to its outer split pair*.

*Proof*. In the SPQR-tree of 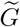, consider first a P-node. Its skeleton consists of multiple parallel edges between two vertices. This structure admits only one possible split pair, which are the endpoints of those parallel edges. Since this is the only connection to the rest of the graph, and all internal paths lie strictly between these two terminals, the directed subgraph inherited from *G* is clearly two-terminal.

Now consider an S-node. Its skeleton is a cycle formed by composing real and virtual edges. Although multiple split pairs may exist in the undirected case, rooting the tree at the Q-node for (*t, s*) selects a unique pair (*x, y*), defined by the virtual edge connecting the S-node to its parent. In the directed version of *G*, let *H* be the subgraph corresponding to this S-node.

Assume for contradiction that *H* is not two-terminal with respect to *x* and *y*. Then there exists a maximal directed path in *H* that begins or ends at a vertex other than *x* or *y*. For example, a path from *x* to *z* where *z* ≠ *y*, and no path exists from *z* to *y* inside *H*. Since *G* is a two-terminal DAG from *s* to *t*, the vertex *z* must still reach *t*, which means it connects outside *H* without going through *y*. This contradicts the fact that *x* and *y* separate *H* from the rest of the graph.

Therefore, both S-nodes and P-nodes define two-terminal subgraphs in *G* with terminals equal to their SPQR-tree split pairs.

A graph *G* is called a series-parallel (SP) graph if its SPQR-tree decomposition contains only *S*-nodes (series compositions), *P* -nodes (parallel compositions), and *Q*-nodes (single edges), and no *R*-nodes. In such graphs, it is straightforward to identify all hetero-positions, as described below.

**Lemma 2**. *Let G be a sequence graph that is series-parallel. Then, hetero-positions occur only at the terminals of the P -nodes in the corresponding SPQR-tree decomposition of G*.

*Proof*. Since *G* is a series-parallel graph, its SPQR-tree contains only *S*-nodes, *P* -nodes, and *Q*-nodes. *Q*-nodes represent single edges and are not relevant for disjoint paths. A *P* -node contains disjoint paths between the same pair of terminals, so those terminals are hetero-positions. *S*-nodes are serial and do not contain disjoint paths between their terminals. Therefore, all hetero-positions in *G* must be at the terminals of *P* -nodes.

Figure 2 shows this idea with an example graph and its SPQR-tree. Part A shows a biconnected sequence graph that is a series-parallel two-terminal DAG with multiple paths from source *s* to sink *t*. The colored vertices mark the terminals of disjoint paths. Part B shows the corresponding SPQR-tree. The *P* -nodes (circles) represent parallel parts of the graph and are colored to match the terminals in Part A. Since only *P* -nodes have disjoint paths between the same pair of vertices, hetero-positions can only appear at their terminals. As shown, the P-nodes and hetero-positions match exactly.

**Figure 2.**
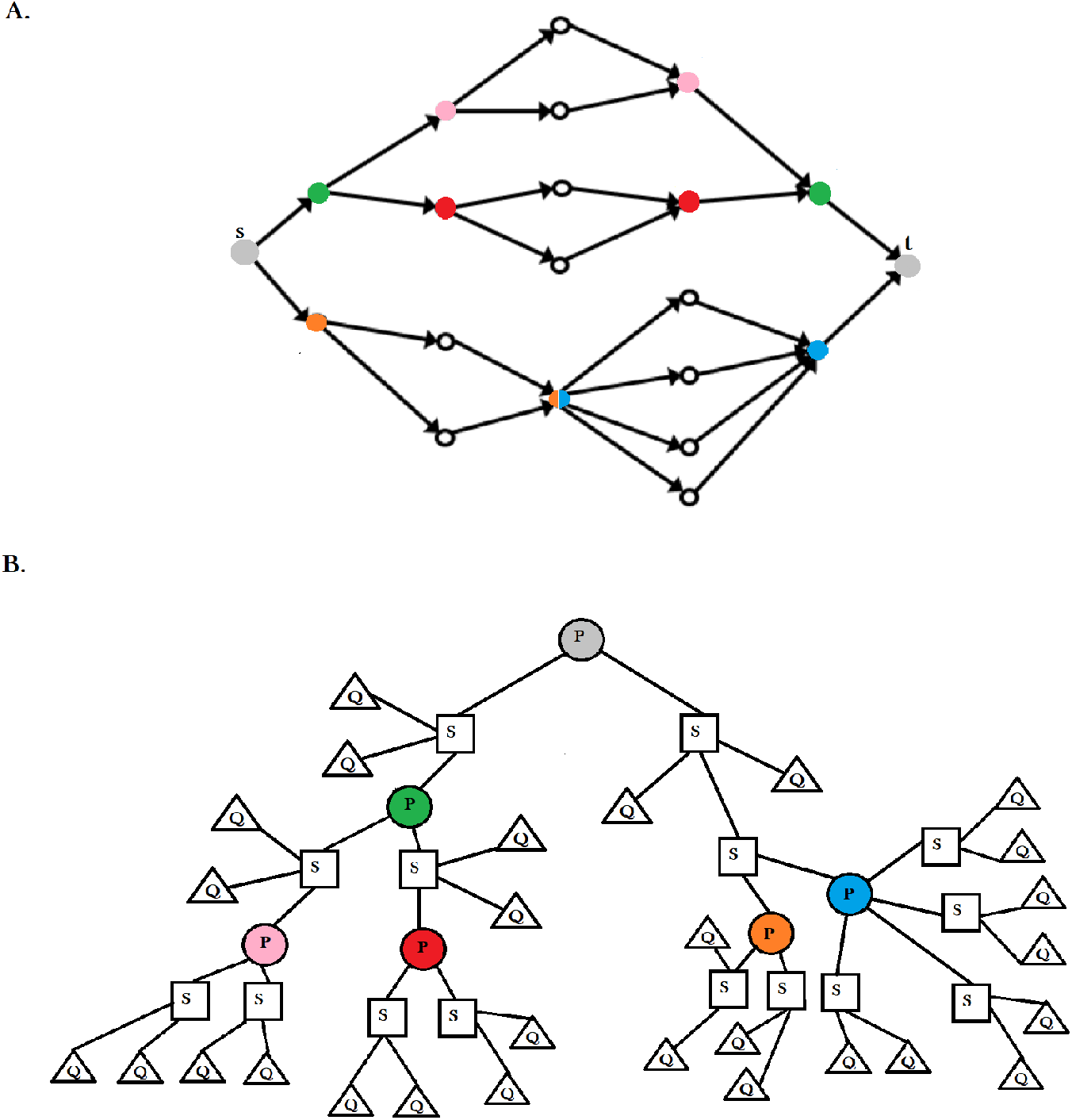
**(A)** A series-parallel sequence graph *G*. Colored vertices indicate pairs of hetero-positions, which are the terminals of parallel structures. **(B)** The SPQR-tree decomposition of the graph in part A. The tree includes *S*-nodes (squares), *P* -nodes (colored circles), and *Q*-nodes (triangles). Each *P* -node is colored to match its corresponding terminals in part A.

Now, let us assume that *G* is non-SP. In the SPQR-tree decomposition of such a graph, at least one R-node must exist. In Valdes’s paper [10], it is proven that non-SP two-terminal digraphs must contain a *W-motif*. A *W-motif* is a directed acyclic graph that is homeomorphic to the graph *W*, as shown in Figure 3. This graph consists of four *main vertices* labeled *u, v, u*^*′*^, and *v*^*′*^, ordered as *u < u*^*′*^ *< v*^*′*^ *< v*. We define the main vertices {*u, v*} as the *terminal vertices* and the main vertices {*u*^*′*^, *v*^*′*^} as the *diagonal vertices*.

**Figure 3.**
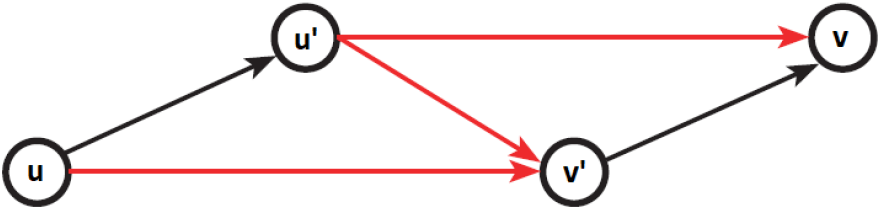
The graph *W*

These vertices are connected in a Z shape, forming two triangles, each consisting of one terminal and both diagonal vertices.

As illustrated in Figure 3, the graph *W* contains three pairs of disjoint paths: one pair between its terminals, (*u → u*^*′*^*→ v, u → v*^*′*^ *→ v*), and two pairs associated with triangles inside the graph, (*u → u*^*′*^*→ v*^*′*^, *u → v*^*′*^) and (*u*^*′*^*→ v*^*′*^ *→ v, u*^*′*^*→ v*).

We then say that each W-motif has three pairs of hetero-positions: one pair related to its terminals and two pairs associated with its triangles.

In Theorem 1, we identify the exact locations of hetero-positions in a two-terminal DAG.

#### Theorem 1.

*In a biconnected two-terminal DAG, any pair of vertices that are shared endpoints of two or more disjoint paths must either be the terminals of a P -node component in the SPQR-tree or among the vertices of a W -motif subgraph*.

*Proof*. Let *G* be a biconnected two-terminal DAG. Suppose there are two vertices {*u, v*} that are the shared endpoints of a set of disjoint paths 𝒫 = {*P*_1_, …, *P*_*k*_}, with *k ≥* 2, where each path shares only the endpoints.

If {*u, v*} is the terminal pair of a *P* -node component, the proof is complete. Otherwise, assume that {*u, v*} is not the terminal pair of any *P* -node. Then we aim to show that {*u, v*} must be among the main vertices of a *W* -motif.

Let *G*_*p*_ be the maximal subgraph of *G* in which every vertex and edge lies on a directed path from *u* to *v*. This subgraph behaves as a two-terminal region, since 𝒫 contains at least two disjoint paths with endpoints *u* and *v*, ensuring its existence.

Since {*u, v*} is not the terminal pair of any *P* -node, *G*_*p*_ is not a *P* -node. Depending on how *G*_*p*_ connects to the rest of *G*, there are two possible cases: either {*u, v*} forms a *separation pair*, or it does not. A separation pair means that removing both *u* and *v*, along with their incident edges, disconnects the graph. In the following, we study each case separately and show that in each case, at least one *W* -motif exists in *G*, with *u* and *v* as two of its main vertices.

**Case 1**. {*u, v*} is a separation pair: Since *G*_*p*_ is not a *P* -node and its terminals {*u, v*} form a separation pair, any two disjoint paths between *u* and *v* must be connected by a path between their internal vertices, meaning a path that does not start or end at *u* or *v*. Otherwise, the disjoint paths would form parallel components, making the region parallel decomposable from the rest of the component, implying that *G*_*p*_ is a *P* -node, contradicting our assumption.

Therefore, for any two disjoint paths *P*_1_ and *P*_2_ between *u* and *v*, there must exist a path *Q* connecting their internal vertices. This connection holds regardless of edge orientation, since SPQR-tree decompositions are based on the undirected structure of the graph.

Let *P*_1_ and *P*_2_ be two disjoint paths connected by a path *Q*. Suppose *Q* connects *u*^*′*^ *∈ P*_1_ to *v*^*′*^ *∈ P*_2_, where *u*^*′*^, *v*^*′*^ ∉ {*u, v*}. If *Q* consists of edges with consistent orientation, forming a directed path (see Figure 4, part I.a), then there exists a *W* -motif in which *u* and *v* are the terminals, and *u*^*′*^ and *v*^*′*^ are the diagonal vertices.

**Figure 4.**
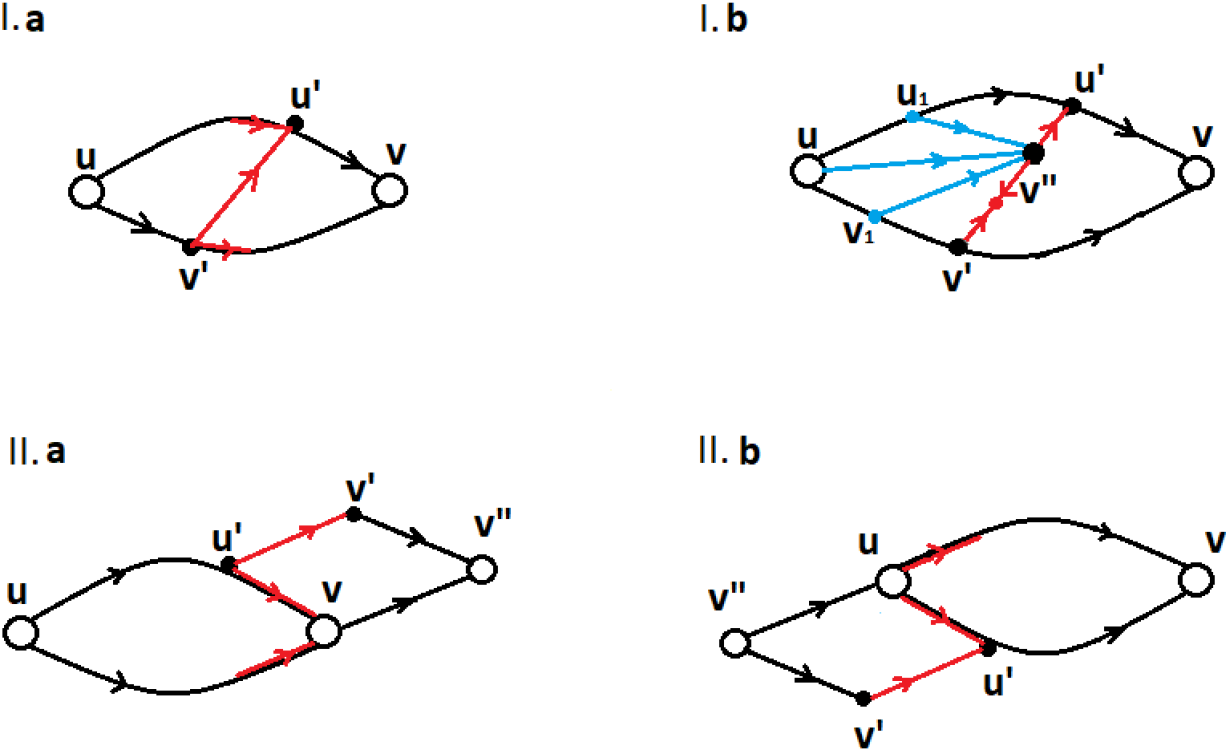
W-motifs in 2 cases of theorem 1

However, if there are no disjoint paths *P*_1_ and *P*_2_ for which a consistently oriented path *Q* connects internal vertices, then let *P*_1_ and *P*_2_ be two disjoint paths connected by a path *Q*, where the length of *Q* is minimized among all such choices of *P*_1_, *P*_2_, and *Q*. By assumption, *Q* is a mixed-direction path connecting *u*^*′*^ *∈ P*_1_ to *v*^*′*^ *∈ P*_2_, as shown in Figure 4, part I.b. Let *v*^*′′*^ be a vertex on *Q* such that the subpath between *u*^*′*^ and *v*^*′′*^ is consistently oriented.

Since *G*_*p*_ is a two-terminal subgraph with terminals {*u, v*}, and we are in the case where {*u, v*} form a separation pair, there must exist a directed path from *u* to *v*^*′′*^, as well as a directed path from *v*^*′′*^ to *v*.

One of these paths already exists, based on the direction between *u*^*′*^ and *v*^*′′*^, which can create a path either from the source *u* to *v*^*′′*^ or from *v*^*′′*^ to the sink *v*. For instance, as shown in Figure 4, part I.b, there is a directed path from *v*^*′′*^ to *v*, which proceeds from *v*^*′′*^ to *u*^*′*^ and then to *v*. We now examine the existence of a directed path from *u* to *v*^*′′*^.

This path must start at *u* and end at *v*^*′′*^, following the edge directions. It may pass through vertices in *P*_1_, *P*_2_, or elsewhere, but it cannot pass through both *P*_1_ and *P*_2_. If it did, it would create a consistently oriented path between internal vertices of *P*_1_ and *P*_2_, which would contradict our assumption that no such paths exist.

So there are three possibilities for this path, as illustrated in Figure 4, part I.b, showing the blue paths on the left. The path can be disjoint from both *P*_1_ and *P*_2_, as shown by the middle disjoint path in the figure. It can share at least one vertex with *P*_1_, as in the upward path passing through *u*_1_ *∈ P*_1_, or it can share at least one vertex with *P*_2_, as in the downward path passing through *v*_1_ *∈ P*_2_.

If there is a directed path from *u*_1_ to *v*^*′′*^, it creates a new directed path *P*_3_ from *u* to *v* through the sequence *u → u*_1_ *→ v*^*′′*^ *→ u*^*′*^ *→ v*. This path *P*_3_ is disjoint from *P*_2_, but there is a connecting path between *P*_3_ and *P*_2_ given by a subpath of *Q*, contradicting the minimality of *Q*.

Similarly, if there is a directed path from *u* to *v*^*′′*^, it creates a new directed path *P*_4_ from *u* to *v* through *u → v*^*′′*^ *→ u*^*′*^ *→ v*. This path *P*_4_ is disjoint from *P*_2_, but there is a connecting subpath of *Q* between *P*_4_ and *P*_2_, again contradicting the minimality assumption.

Finally, if there exists a directed path from *v*_1_ to *v*^*′′*^, it yields a consistently oriented path connecting *P*_1_ to *P*_2_ via the sequence *v*_1_ *→ v*^*′′*^ *→ u*^*′*^. This contradicts our assumption that no such consistently oriented path exists between *P*_1_ and *P*_2_ through their internal vertices.

Since all possible configurations lead to a contradiction, our assumption must be false. Therefore, there must exist at least two disjoint paths *P*_1_ and *P*_2_ with a consistently oriented path *Q* connecting their internal vertices.

As shown earlier, such a configuration gives rise to a *W* -motif with terminals *u* and *v*, as illustrated in Figure 4, part I.a.

**Case 2**. {*u, v*} is not a separation pair: In this case, there must be at least one connection from an internal vertex of *G*_*p*_ to a vertex outside this subgraph in *G*.

Suppose there is a vertex *u*^*′*^ *∈ G*_*p*_ and a vertex *v*^*′*^*∈ G* − *G*_*p*_. If *u*^*′*^ *< v*^*′*^, then since *G* is a two-terminal directed graph, there must exist a directed path from *u*^*′*^ to *v*^*′*^.

Also, there must exist a vertex *v*^*′′*^ such that both *v* and *v*^*′*^ are connected to *v*^*′′*^, where *v*^*′′*^ is the closest such vertex. This configuration forms a *W* -motif in which *u* and *v*^*′′*^ are the terminal vertices, and *u*^*′*^ and *v* are the diagonals, as illustrated in Figure 4, part II.a.

A similar argument applies when *v*^*′*^ *< u*^*′*^. In that case, we obtain a *W* -motif where *v*^*′′*^ and *v* are the terminal vertices, and *u*^*′*^ and *u* are the diagonal vertices, as shown in Figure 4, part II.b.

Note that *v*^*′′*^ may be the same as *v*^*′*^, but it cannot be equal to *v* or *u*, as that would imply *v*^*′*^ lies on a path from *u* to *v*, contradicting the assumption that *v*^*′*^ ∉ *G*_*p*_.

Thus, in this case, a *W* -motif must exist in which one of *u* or *v* is a terminal vertex, and the other appears as a diagonal vertex.

We have shown that if {*u, v*} are not the terminals of a *P* node, then in both cases, whether {*u, v*} form a separation pair or not, they must be among the main vertices of a *W* -motif. In Case 1, *u* and *v* appear as the terminal vertices of the motif. In Case 2, *u* and *v* appear as either terminal or diagonal vertices of a *W* motif.

Therefore, we conclude that in a biconnected two-terminal DAG, any pair of vertices that are the shared endpoints of two or more disjoint paths must either be the terminals of a *P* node or among the main vertices of a *W* motif. This completes the proof.

The question now is: *How can we identify W-motifs in a non-SP sequence graph?*

To do this, we briefly introduce the concept of the *complexity graph C*(*G*), originally proposed by Bein et al. [11]. This structure provides a formal measure of how far a two-terminal DAG is from being series-parallel, and is central to our identification of W-motifs. Our approach leverages *C*(*G*) as a tool to localize non-series-parallel substructures, those embedded within R-nodes of the SPQR decomposition.

#### Definition 1

(Complexity Graph [11]). *Let G be a two-terminal directed acyclic graph (DAG) with unique source s and sink t. The* complexity graph *C*(*G*) *is a directed graph where an edge* (*x, y*) *∈ C*(*G*) *if and only if:*

- *x < y, i*.*e*., *there exists a directed path from x to y in G*,
- *There is no vertex z ≥ x that properly dominates y, and*
- *There is no vertex z ≤ y that properly reverse-dominates x*.

*A vertex v is said to* dominate *a vertex w if every path from the source s to w passes through v. Conversely, a vertex w* reverse-dominates *v if every path from v to the sink t passes through w. Dominance must be* proper, *meaning v* ≠ *w*.

This definition captures the structural complexity of *G*: each edge in *C*(*G*) corresponds to the diagonal path of a W-motif, a forbidden configuration in series-parallel DAGs. Therefore, *C*(*G*) has no edges if and only if *G* is series-parallel.

For example, in Figure 1, part A, the graph *G* consists of two biconnected components, each being a two-terminal DAG. One component, with terminals {*a, c*}, is SP, and hence its complexity graph *C*(*G*) has no edges, as no pair of vertices satisfies the complexity conditions. The other component, with terminals {*c, b*}, is non-SP, and its complexity graph *C*(*G*) contains precisely one edge from vertex *k* to vertex *d*. These are the only two vertices satisfying the complexity conditions: no vertex *z ≥ k* properly dominates *d*, and no vertex *z ≤ d* properly reverse-dominates *k*.

In the following lemma, we formally prove the relationship between edges in *C*(*G*) and the diagonal vertices of W-motifs in non-SP DAGs.

#### Lemma 3.

*Let G be a biconnected two-terminal DAG. For every pair of vertices x and y in G*, (*x, y*) *∈ C*(*G*) *if and only if there is a W-motif in G with* {*x, y*} *as the diagonal vertices*.

*Proof. If part:* Let *x* and *y* be vertices in *G*, and suppose (*x, y*) *∈ C*(*G*). Then *x < y*, which means there exists a path *P*_*xy*_ from *x* to *y*. Based on the definition of *C*(*G*), there is no vertex on *P*_*xy*_ that properly dominates *y*, meaning there must be a path *P*_*sy*_ from the source vertex *s* to *y*, where the only common vertex between *P*_*xy*_ and *P*_*sy*_ is *y*. Similarly, since there is no vertex on *P*_*xy*_ that reverse-dominates *x*, there must be a path *P*_*xt*_ from *x* to the sink vertex *t*, where the only common vertex between *P*_*xy*_ and *P*_*xt*_ is *x*. As shown in Figure 5, the union of paths *P*_*sy*_, *P*_*xy*_, and *P*_*xt*_ forms a Z-shape, which corresponds to a W-motif in *G*. This means that there exists a W-motif with diagonal vertices {*x, y*}.

**Figure 5.**
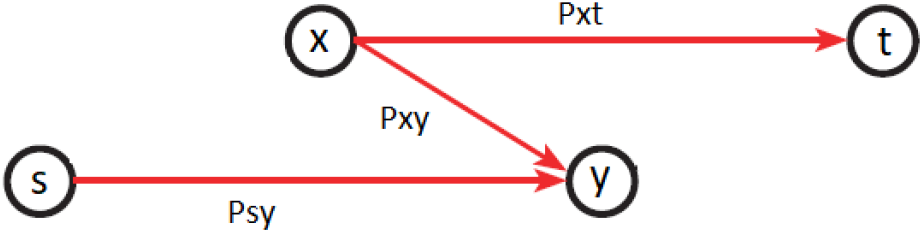
Z-shape in Lemma 2

*Only if part:* Let *P*_*xy*_ be the diagonal path of a W-motif. Based on the Z-shape of each W-motif, it is obvious that *x < y*, and there is no vertex *z ≥ x* that properly dominates *y*, because of the existence of a path *P*_*sy*_ from *s* to *y* with no common vertex with *P*_*xy*_. Similarly, there is no vertex *z ≤ y* that properly reverse-dominates *x*, because of the existence of a path *P*_*xt*_ from *x* to *t* with no common vertex with *P*_*xy*_. Therefore, (*x, y*) *∈ C*(*G*).

### 3 Diploid Graph Construction

We now give an algorithm to compute all the W-motifs by constructing the *SPQRW-tree*, an extension of the SPQR-tree that includes *W-nodes*. These nodes are attached to R-nodes, and each represents a W-motif whose diagonal path lies within the corresponding R-node. Each R-node has at least one W-node as a direct child.

Our algorithm consists of three steps, using efficient existing techniques to analyze the graph and identify its decomposition components. The procedure is described below and formally presented in Algorithm 2.

#### Algorithm 2

Constructing SPQRW-Tree for Two-terminal DAGs

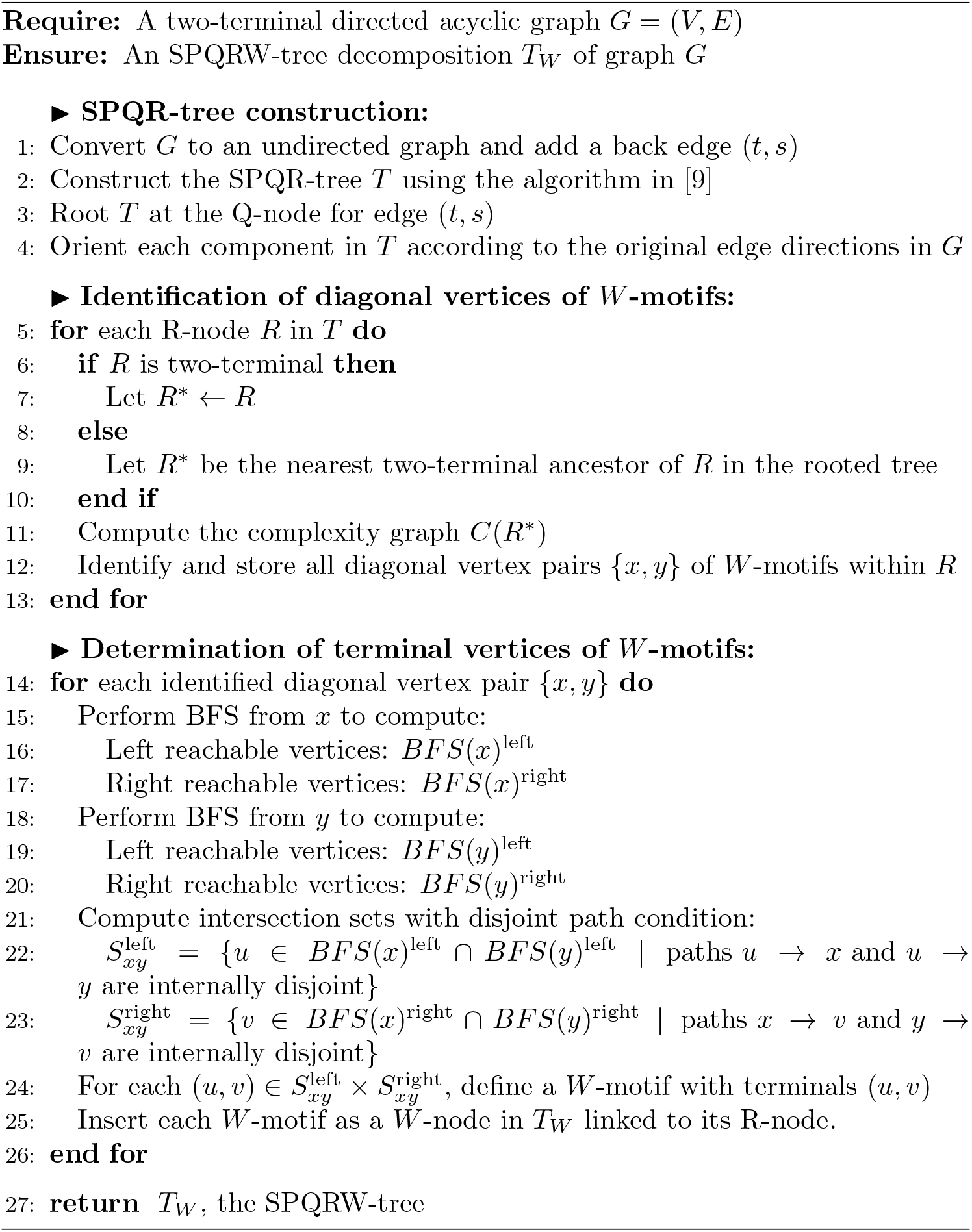

**Step 1**. Convert *G* into 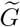 by making it undirected and adding a back edge. Construct its SPQR-tree rooted at the Q-node for that edge, then reassign directions using the original orientations in *G*.

**Step 2**. To enumerate the diagonal vertices of *W* -motifs, as described in Lemma 2, we compute the complexity graph of *G*, denoted *C*(*G*). An efficient polynomial-time algorithm with complexity *O*(*n*^2.5^) is provided in [11]. To reduce runtime, we apply it only to relevant components of the SPQR-tree rather than the full graph. Since the tree is already rooted, we define *R*^*\**^ for each R-node as its nearest two-terminal ancestor (possibly the node itself). We then compute *C*(*R*^*\**^) and extract all diagonal vertex pairs corresponding to *W* -motifs.

**Step 3**. After identifying the diagonal vertices of the *W* -motifs, we determine their terminal vertices using Breadth-First Search (BFS). For each pair {*x, y*}, we run BFS from both *x* and *y*. We then collect all vertices that are reachable from both *x* and *y* through paths that do not share any internal vertex, except for the target itself. Each such vertex defines a possible terminal point of a *W* -motif. We allow all valid terminal vertices for each diagonal pair, since a single diagonal pair may correspond to multiple terminal pairs, and thus to multiple *W* -nodes. For each motif, we consider all combinations of one predecessor and one successor of the diagonal pair to define terminal vertex pairs. The overall time complexity of this step is *O*(*n*^2^), since we perform two BFS traversals for each diagonal pair.

Based on the described steps, the time complexity of the algorithm for constructing an SPQRW-tree is *O*(*n*^2.5^), where *n* is the number of vertices in the largest *R*^*\**^ component in the SPQR-tree decomposition. However, we expect it to be significantly faster than the worst case bound in practice, since pangenome graphs tend to be largely series-parallel, and the worst case run time of this algorithm is due to R-nodes. After constructing the SPQRW-tree using Algorithm 2, the decomposition includes all necessary *P* -nodes and *W* -nodes, which serve as the structural patterns for extending the sequence graph to obtain a diploid sequence graph.

#### Algorithm 3

Diploid Graph Construction via SPQRW-tree

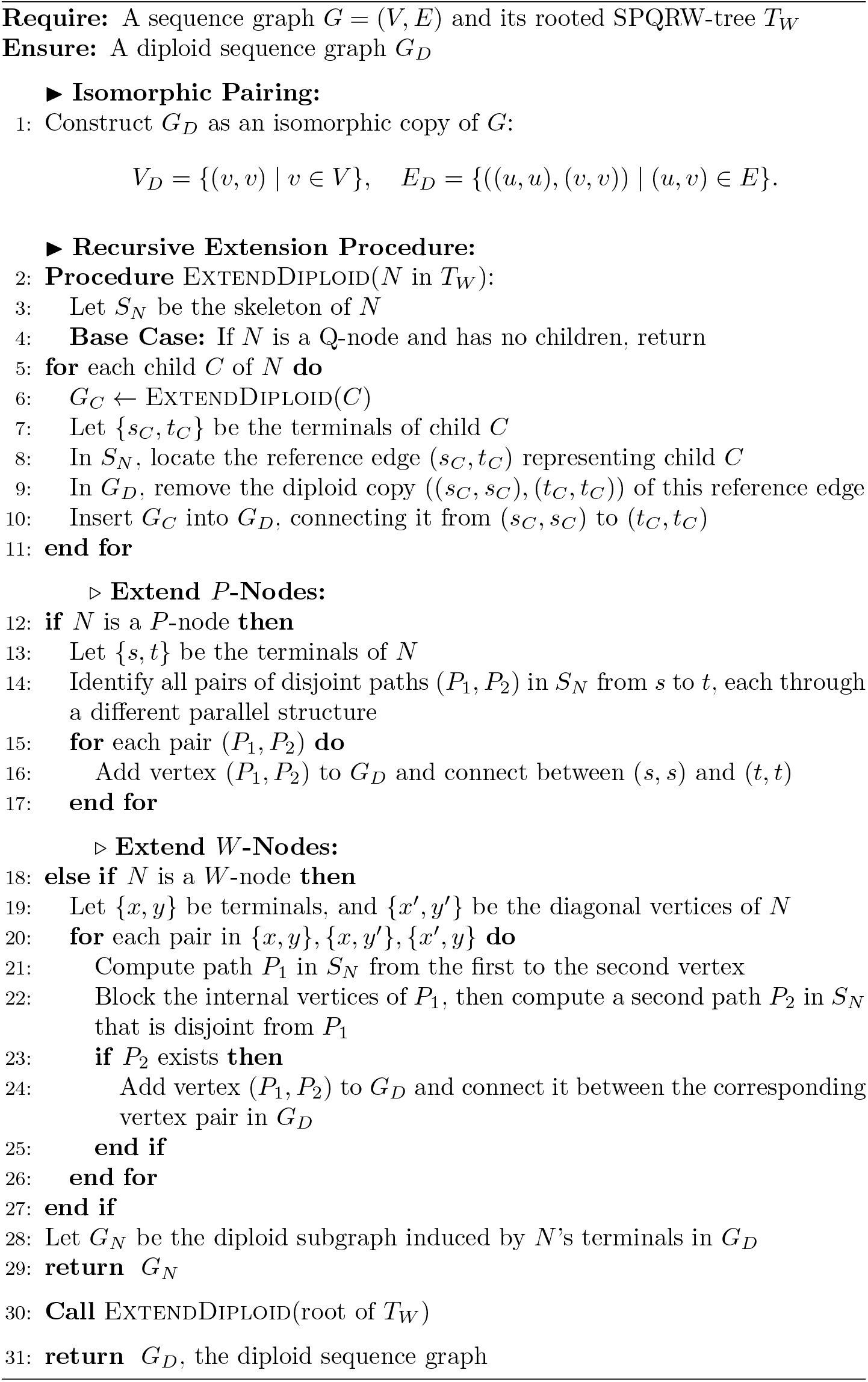

In Algorithm 3, we present a recursive procedure for constructing the diploid sequence graph based on the SPQRW-tree decomposition. This algorithm is a detailed version of the general framework in Algorithm 1, where hetero-positions are identified based on the graph’s structural decomposition.

The construction starts by creating an isomorphic copy of the input sequence graph. It then recursively traverses the rooted SPQRW-tree in post-order. This ensures that each component is extended only after its children have been processed, preserving the hierarchical structure of the decomposition.

New vertices are added only at *P* - and *W* -nodes. Other node types are visited but skipped during extension. Figure 6 illustrates how relevant node types contribute to the diploid structure. While all components are copied isomorphically, only *P* - and *W* -nodes generate variant-specific vertices.

**Figure 6.**
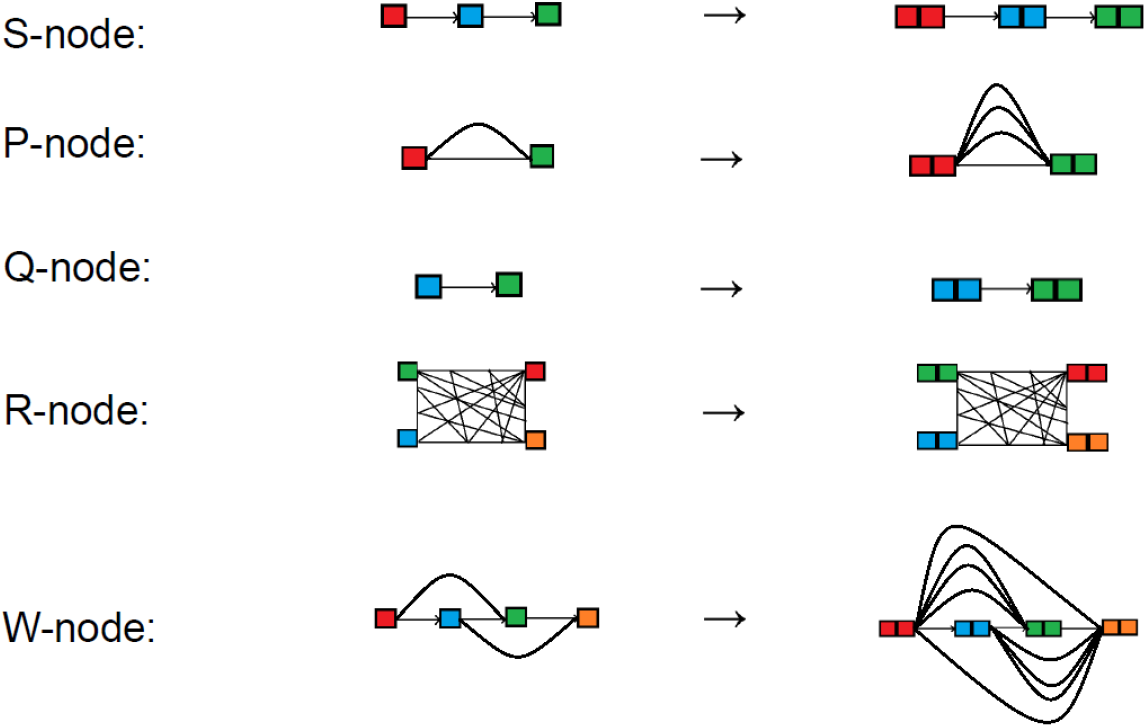
Extension Rules for Components of the SPQRW-tree

The validity of Algorithm 3 follows directly from Algorithm 1, as established in Proposition 1. In Proposition 2, we further analyze the worst-case time complexity of Algorithm 3 when the SPQRW-tree is provided as input, demonstrating its efficiency and scalability in practice.

#### Proposition 2

(Speed of Algorithm 3). *The worst-case time complexity of Algorithm 3 is:*

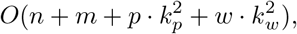

*where n and m are the number of vertices and edges in the input graph G, and p and w are the number of P -nodes and W -nodes in the SPQRW-tree T*_*W*_. *The term k*_*p*_ *is the maximum number of disjoint path pairs between the terminals of any P -node. The term k*_*w*_ *is the maximum number of disjoint path pairs evaluated in any W -node, each connecting either two terminals or a terminal and a diagonal vertex*.

*Proof*. The algorithm consists of three main steps:

Step 1: Constructing *G*_*D*_ as an isomorphic copy of *G* involves duplicating all vertices and edges, which takes: *O*(*n* + *m*).

Step 2: For each *P* -node, finding all paths between *s* and *t* using DFS takes 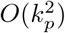. Pairing two disjoint paths from at most *k*_*p*_ parallel paths takes *O*(*k*^2^). With *p P* -nodes, this contributes: 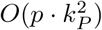.

Step 3: For each *W* -node, identifying valid path pairs using DFS takes *O*(*m*). Pairing two disjoint paths from at most *k*_*w*_ possible paths takes 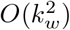. With *w W* -nodes, this contributes: 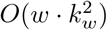.

Summing these contributions, the overall time complexity is:

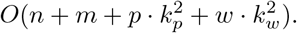

## Discussion

Accurately representing diploid genomes is a fundamental challenge in modern genomics, especially as the field moves toward graph-based models to capture population-scale variation. While sequence and variation graphs have proven effective for representing haplotypes, extending these models to fully capture diploid complexity, including structural variation and recombination, has not been sufficiently explored, either computationally or conceptually.

Our work addresses this gap by identifying the minimal structural features required for principled and efficient diploid graph construction. Rather than pairing all maximal haplotype paths, we introduce a framework that targets disjoint path regions corresponding to true heterozygosity. To detect these regions systematically, we build on the classical SPQR-tree decomposition and extend it with a new structure, the SPQRW-tree. The key structural patterns identified through this process, namely P-nodes and W-nodes, act as anchors for graph extension and ensure that all relevant diploid sequences are represented without redundancy.

Beyond topological completeness, our model is designed to support probabilistic modeling. A key strength of our approach is that it provides a clean topological foundation for future probabilistic models. In our future work, we will use the diploid sequence graph to structure the state space of a Hidden Markov Model (HMM) for use in diplotype inference. This will enable modeling recombination and haplotype-specific variation in a probabilistic framework, while maintaining alignment with the underlying graph structure.

This model offers a robust foundation for diploid modeling and inference, enabling more accurate and biologically meaningful analysis of diploid genomes. It also supports a range of applications, including haplotype sampling, haplotype-aware genome assembly, and structural variation analysis in pangenomics.

